# A pilot computer-vision model trained to identify slide-mounted scale insect pests from the family Rhizoecidae

**DOI:** 10.64898/2026.10.16.759574

**Authors:** Azizah Conerly, Benjamin Rosen, Scott A. Schneider

## Abstract

The emerging field of image classification using computer vision has yet to address a rather challenging group of insect pests—soft-bodied insects that are preserved on microscope slides. This includes many notoriously invasive herbivores that have a significant impact on agriculture, such as members of the hemipteran suborder Sternorrhyncha. Using root mealybugs (Hemiptera: Rhizoecidae) as a test case, we gathered high-resolution extended depth of field images from museum specimens and trained a convolutional neural network (CNN) to identify and distinguish among 16 root mealybug species that are encountered as agricultural pests in plant quarantine or greenhouses. A pretrained EfficientNetV2-M model was trained using BioEncoder. Our dataset contained 1,607 images, with 41 to 239 images per category. Performance was improved through the adoption of augmentation techniques. Upon completion of training, the model achieved macro precision, recall, and F1 scores of 80.1%, 79.3%, and 79.1%, respectively. The model demonstrated classification metrics >= 80% for seven species. Gradient-weighted Class Activation Mapping (Grad-CAM) was used to visualize the morphological features associated with the test-image classification. Grad-CAM images highlighted meaningful diagnostic features to distinguish among species rather than focusing on uninformative visual artifacts or specimen damage, demonstrating the model’s utility for identification purposes. The results suggest that further development of the model should incorporate at least 200 training images per species with a minimum resolution of 600×600 pixels. The model was designed to be adaptable; images can be added to improve its performance, and more species can be incorporated as specimens become available, expanding on the taxonomic breadth of its application. Computer-vision identification tools have the potential to enhance rapid response capabilities in plant quarantine at ports-of-entry in the United States and globally.

## Introduction

Scale insects (Hemiptera: Coccomorpha) are frequently encountered in plant quarantine at global ports-of-entry and many species have become notorious pests of agriculture, horticulture, and forestry. Non-native scale insects are responsible for substantial economic losses due to damage and management costs. In the United States alone, scale insects are estimated to cost agriculture and consumers minimally $4–8 billion each year [1]. The rapid, accurate identification of insects that are intercepted in plant quarantine is a critical preventative step to impede the spread of invasive species and limit agricultural losses while fostering trade. Plant quarantine serves as an important and effective barrier to the spread of invasive species [2, 3]. But, approximately one new non-native scale insect species establishes in the United States each year [1, 4], indicating there is room to improve on preventative measures, such as developing more comprehensive and more accessible identification tools that support quarantine and general pest identification efforts.

Scale insects can be difficult to identify, requiring specialized training and expertise [5, 6]. For accurate identification and preservation, a scale insect specimen is cleared of its body contents and the remaining cuticle is mounted on a microscope slide [7]. Scale insect taxonomy is based upon adult females, the longest lived and most reliably identifiable life stage. Challenges relating to the identification of scale insects often involve their small size, sexual dimorphism, neoteny (of adult females), microscopic diagnostic features, life-history characteristics, and cryptic species diversity [5, 8]. The challenge of identifying scale insects is exacerbated by the taxonomic impediment [9–11], inadequate investment in the field of taxonomy and loss of expertise through attrition. The number of experts trained in their identification is diminishing, which presents an important practical problem, especially for plant quarantine. This loss of expertise coincides with extensive and increasing international trade in agricultural and horticultural goods to the U.S. [1, 12–15] and notably high propagule pressure of scale insects [16].

Computer-vision models offer a potential avenue to support taxonomic experts and non-experts alike in the identification of slide-mounted scale insects. Historically, scale insect systematists have been early adopters of alternative diagnostic tools [6], including the use of ScaleNet [17] and matrix-based interactive keys [18, 19]. To explore the potential for computer-vision models to identify challenging, minute slide-mounted insects we trained a convolutional neural network (CNN) to identify a set of 16 pest species. This pilot project focuses on root mealybugs (Hemiptera: Rhizoecidae) as a test case. Root mealybugs are a relatively understudied family of scale insects that may impact agriculture more than appreciated due to their subterranean habit [20–24]. They can be difficult to identify to species, even for experts. Drawing upon interception records from the USDA’s Animal and Plant Health Inspection Service (APHIS), about 50% of root mealybug interceptions have been identified to species level, with the remaining ∼50% identified to genus or family level only (unpublished data). This is comparable to metrics for the three largest scale families comprising most agricultural scale insect pests in the U.S.—Coccidae, Diaspididae, and Pseudococcidae [1, 4, 25–27]. A computer-vision model that can reliably identify rhizoecid pests, as an example, could benefit regulatory agencies by offering (i) fast, accurate diagnosis to species-level for a greater percentage of quarantine seizures, (ii) efficient time allocation for overextended taxonomic experts [28], and (iii) more refined data on host-associations, transmission of species, and potential invasion pathways. The availability of such tools could enhance national biosecurity by improving practices for invasive species prevention and open new avenues for cutting-edge research on morphology and systematics. These models can also be expanded and re-trained to include broader taxonomic representation as more data become available.

Slide-mounted insect specimens are limited to a 2-dimensional view, and the fixed body orientation of a mounted specimen is preparation-dependent. Across series of specimens, inconsistencies will occur in the clearing of body contents, body orientation, appendage orientation, specimen damage, the color and intensity of staining, and the presence of various background artifacts in the mounting medium (Fig 1). Thus, a CNN model may have to learn a wider range of distinctive features for accurate identification [28] and learn to disregard image artifacts. Similar challenges have applied to models trained in the identification of diatoms [29], protozoan intestinal parasites [30], and fungal genera [31]. For insects, many CNNs have been trained on taxa with sclerotized exoskeletons and rigid structures, which allows for greater consistency in orientation of views [28, 32–34] and reduces the imaging burden required per class, especially when combined with transfer learning [34]. In this respect, soft-bodied insects on glass slides differ. The approach to model training must address challenges inherent to slide-mounted collections and may require larger image training sets.

**Fig 1.**
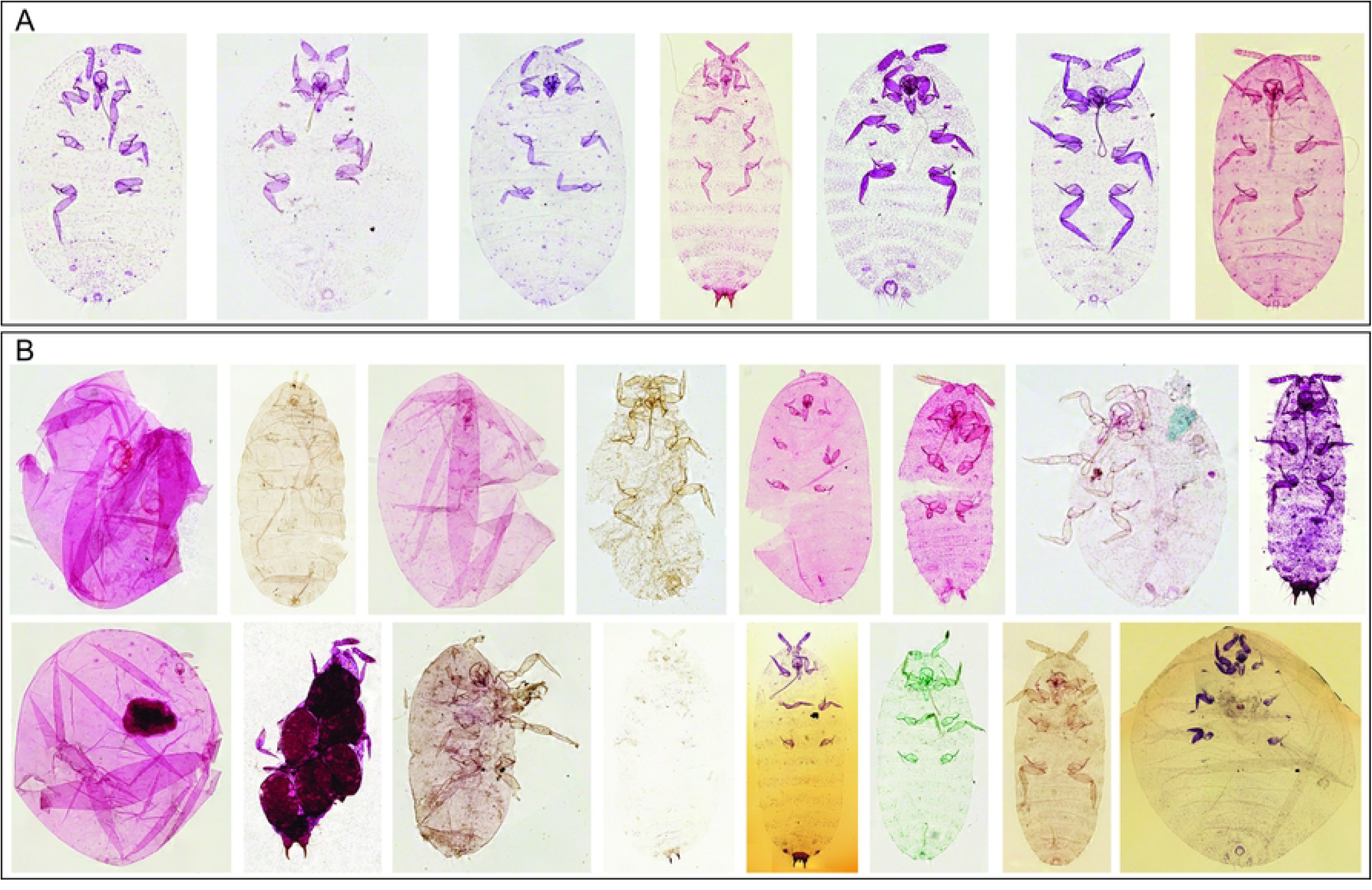
Preparation-dependent inconsistencies in slide-mounted specimens. (A) Representatives of high-quality, intact specimens in standard orientation; (B) representatives with inconsistencies inherent to slide-mounted collections, including variations in orientation, specimen damage, clearing of internal contents, staining, coloration, and presence of background artifacts.

The aims of this pilot study were to determine limitations and best practices for CNN model training on slide-mounted insect specimens, including selecting the best pre-trained model, determining the ideal minimum image training set size, minimum image resolution, and the most suitable suite of augmentations. The model demonstrated classification metrics >= 80% for seven species, with success being driven primarily by the number of images available for training but also by how distinctive certain classes were morphologically, as assessed using Gradient-weighted Class Activation Mapping (Grad-CAM). The model achieved macro precision, recall, and F1 scores of 80.1%, 79.3%, and 79.1%, respectively.

## Materials and Methods

### Whole-Drawer Slide Imaging

We targeted root mealybug species that had 40+ images available for inclusion in the study, with a range of 41–239 images per species (Table 1). Study specimens were drawn primarily from the U.S. National Entomological Collection (USNM), but we also borrowed material from the California Department of Food and Agriculture (CDFA) and Florida State Collection of Arthropods (FSCA). Each collection is largely representative of species that have significance to agriculture or horticulture; the target species are often collected from quarantine interceptions, greenhouses, nurseries, or other agricultural environments (fields, plantations, etc.) [17, 35].

**Table 1.** Species selected for inclusion in the CNN model with the number of images for the training (80%), validation (10%) and testing (10%) datasets listed.

| Species | Training | Validation | Testing |
| --- | --- | --- | --- |
| *Geococcus coffeae* Green | 185 | 24 | 24 |
| *Rhizoecus americanus* (Hambleton) | 98 | 13 | 13 |
| *Rhizoecus amorphophalli* Betrem | 31 | 5 | 5 |
| *Rhizoecus cacticans* (Hambleton) | 49 | 7 | 7 |
| *Rhizoecus coffeae* Laing | 58 | 8 | 8 |
| *Rhizoecus colombiensis* Ramos & Caballero | 47 | 7 | 7 |
| *Rhizoecus dianthi* Green | 82 | 11 | 11 |
| *Rhizoecus distinctus* (Hambleton) | 37 | 5 | 5 |
| *Rhizoecus falcifer* Kunckel d'Herculaïs | 119 | 16 | 16 |
| *Rhizoecus floridanus* Hambleton | 91 | 12 | 12 |
| *Rhizoecus leucosomus* (Cockerell) | 50 | 7 | 7 |
| *Ripersiella gracilis* (McKenzie) | 60 | 8 | 8 |
| *Ripersiella hibisci* (Kawai & Takagi) | 59 | 8 | 8 |
| *Ripersiella kelloggi* Ehrhorn & Cockerell | 191 | 24 | 24 |
| *Ripersiella kondonis* (Kuwana) | 45 | 6 | 6 |
| *Ripersiella palestineae* Hambleton | 65 | 9 | 9 |

We digitized and captured high-resolution images for the full USNM collection of Rhizoecidae plus material on loan from CDFA and FSCA. Each USNM slide received a unique accession number (ex: USNMENT02074062) and we compiled associated label metadata into a data matrix. Images were captured using a Glissando 20SL (Objective Imaging, Cambridge, UK) under an ultra-high-resolution Plan Apo 20x /075 n.a. objective. This semi-automated imaging system acquires a picture of the slide and scans a user-defined section of the coverslip region containing specimens at a sensor image resolution of 0.5 μm. Scanning is performed in tiles acquiring Z-stack images with a step size of 0.8 μm and a typical scanning range of 70 μm (typically 88 Z-stacks per tile). Tiles are automatically stitched and compiled into an extended depth of field (EDF) image as output. Integrated EDF imaging allowed for rapid, high-resolution capture with both dorsal and ventral features displayed in one image. For slides with multiple root mealybugs, we segregated specimens into individual image files and assigned each file a sequential identifier tied to the accession (ex: USNMENT02074062_1) (Fig 2). We assigned unique identifiers to images captured from loaned material by using the collection acronym (ex: CDFA) and existing identification numbers or dates from the label data. We edited images using *Inselect* [36] but used manual segregation boxes instead of automated specimen segregation, which often focused on debris beneath the coverslips. For each image file, specimen orientation was standardized to the extent possible along the anteroposterior axis using Microsoft Photos (Microsoft Corporation, Redmond, WA).

**Fig 2.**
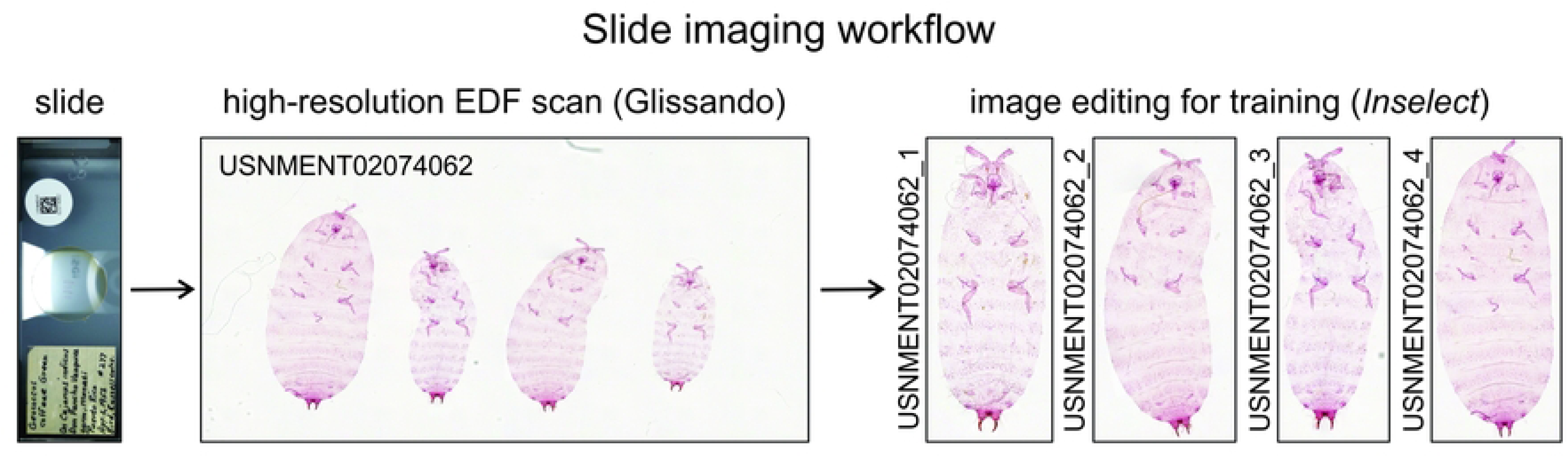
Whole-drawer slide imaging workflow. The process included cataloguing and digitizing the full USNM collection of Rhizoecidae, scanning slide images, scanning specimens under high resolution in a user-defined window, parsing specimens into individual image files, and editing the anteroposterior orientation of specimens for greatest possible consistency in model training.

### Training and Testing Dataset

Fig. 2 shows some representative specimens included in model training and the workflow for acquiring images. Some specimens have shifted body or appendage orientations, variations in clearing or staining, or specimen damage that deviated from the standard positioning. Such inconsistencies are inherent to slide-mounted collections. Images containing immature or male specimens were filtered from the training dataset. We also excluded specimens that were likely to be too damaged to make a proper identification, as determined by the taxonomic expert. However, some immature or damaged specimens were retained in testing datasets to assess limitations of the model for identification (S1 Table).

Due to an imbalanced distribution of the imaged species, we used a stratified random split to preserve the species proportions in the subsets. The subsets include the training (80%), validation (10%), and test (10%) sets.

A randomly selected set of 30 species that were not included in model training served as an additional ‘outgroup’ dataset for testing, using one image per species. The purpose of testing on outgroup species was to assess how the model assigns confidence to correct predictions of species included vs. incorrect predictions for confamilial species excluded from model training. Outgroup species were selected from among the remaining digitized collections, including three species each of *Coccidella* and *Geococcus*, and 12 species each of *Rhizoecus* and *Ripersiella* (S1 Table).

### Data Preprocessing

During preliminary training, images were resized to squares of 480×480 and 600×600 pixels. Based on the initial investigation an image size of 600×600 pixels was selected. This method may distort or skew the specimen, introducing bias or information loss that affects training. As this was a pilot study, the square resizing method was deemed appropriate for the initial implementation. Future studies may include stages to segment the specimen from the background, preserving the aspect ratio.

The dataset used in this work contains 1,607 images. To expand the dataset, data augmentations were systematically applied. Transformations were selected based on their improvements in model robustness on the test dataset. The following transformations were included in the final model: CLAHE, HorizontalFlip, RandomRotate90, and HueSaturationValue.

### Model training

We utilized a supervised deep learning approach. A training setup was developed, including a pipeline for training and testing in Python, using BioEncoder [37] and TorchMetrics. The Python scripts and a list of all used libraries and their versions can be found in S1 Appendix.

Transfer learning was employed to leverage a CNN previously trained on a large image dataset for a generic image classification task [38]. This allowed fewer images to be used for training. The EfficientNet model architecture was selected for its performance on classifying species using other insect datasets [39–43]. The modern version of EfficientNet, EfficientNetV2-M trained on ImageNet-21k images and fine-tuned on ImageNet-1k images, was selected for training due to its improvements in training speed, parameter efficiency, and increased accuracy [44]. The first stage of training used the pretrained EfficientNetV2-M model architecture without a classification head for feature extraction. The model was trained for 100 epochs. Exponential moving average was used to stabilize training. The Stochastic Gradient Descent optimizer with CosineAnnealingLR learning rate scheduler was employed to optimize the learning rate during model training. We used the SupCon loss function. Stochastic weight averaging (SWA) was applied to the top three performing model weights to improve generalization of the model. The most effective learning rate for the second stage was automatically selected using BioEncoder, ensuring the pipeline required no human intervention. The second stage was trained for 30 epochs with a classification head and used a cross-entropy loss function with label smoothing. We again applied SWA to the top three performing model weights to improve the generalizability of the model. Details on training parameters and augmentations can be found in S2 Appendix. All computations were completed using an NVIDIA V100 GPU.

### Evaluation

Classification performance was evaluated using precision, recall, and F1-score on the test set without the outgroup. Accuracy was not considered, as it may become skewed when used as the defining metric for imbalanced datasets. Metrics were calculated using TorchMetrics. To ensure equal representation for the multi-class imbalanced dataset, macro-averaged metrics were used. Performance scores were calculated for each class independently, and the final score was calculated as the unweighted mean. Precision and recall were calculated for each class *n*, then the macro scores were calculated as follows:

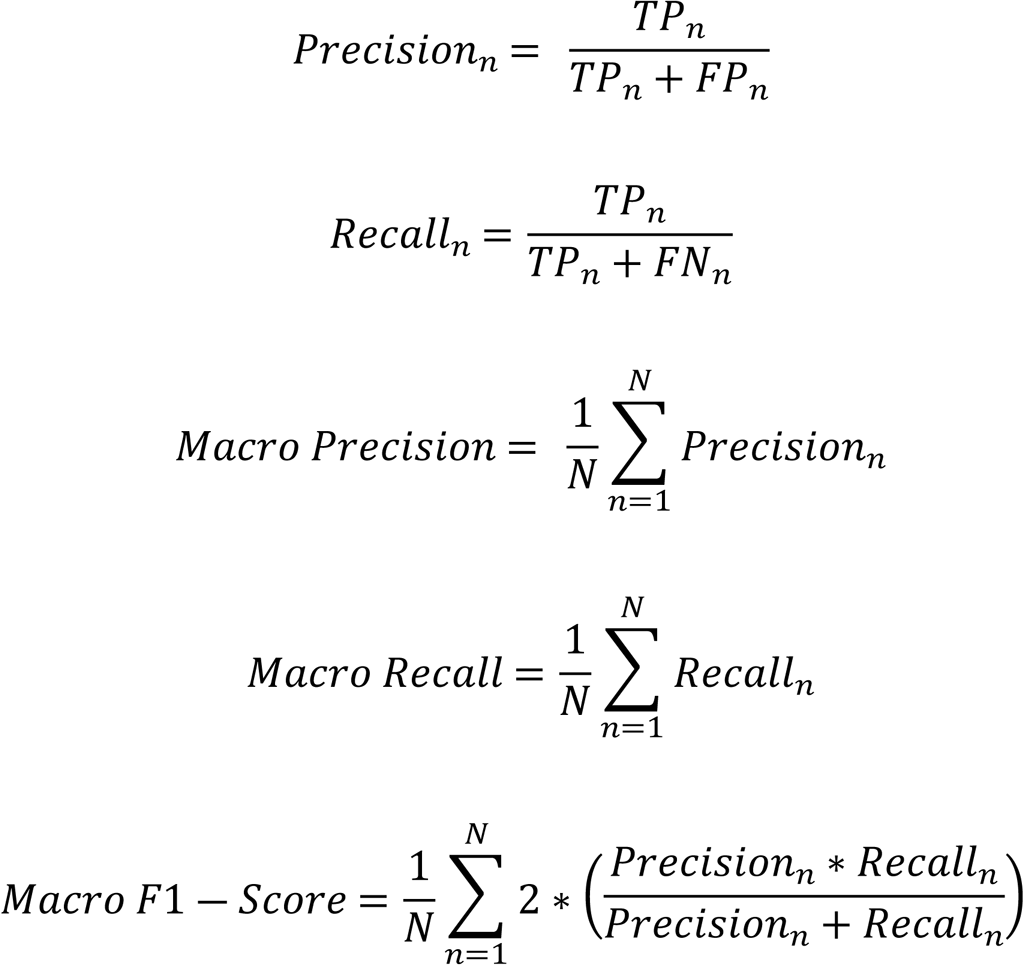

To ensure the model selected morphologically relevant features we employed Grad-CAM with BioEncoder. Grad-CAM is a class activation map that identifies the pixels that most affect the final classification [45].

A Kruskal-Wallis test was used to assess differences in confidence scores assigned among correctly predicted specimens, incorrectly predicted specimens, and the outgroup set of species tested. The purpose was to determine if the model is significantly more confident in its predictions on average when they are correct. To determine if the result is significant, an alpha of 0.05 was selected. To determine which test group differed, a post-hoc Wilcoxon rank-sum test with continuity correction was performed with an alpha of 0.05. Analyses were conducted in R using the packages *dplyr* [46], *ggplot2* [47], *ggpubr* [48], and *readr* [49]; R code is included in S3 Appendix.

## Results

### Whole-drawer Slide Imaging

The full USNM collection of Rhizoecidae was imaged excluding slides with physical damage, non-standard proportions, or mounting techniques that yield a non-centered coverslip [50]. Image datasets were supplemented with loans from CDFA and FSCA (S2 Table). The combined imaging effort produced 2,630 individual specimen images from 1,406 slides. Time required to complete a 20-slide imaging run using the Glissando slide scanner was dependent on specimen size and number of specimens per slide (i.e., the scanning window size), with a mean runtime of 89.51 minutes (SD = 23.39), which equates to about 4.5 minutes per specimen. Image editing was conducted in parallel with the imaging workflow and added about one minute per slide. Overall, imaging and editing are estimated to have totaled about 221 hours of effort. Thus, this work could be achieved by one person within a period of about six weeks minimally under a standard 40-hour work schedule if imaging and editing were the only tasks performed.

### Classifier Performance

To select the minimum image resolution, an initial base model was trained with image resizing to 480×480 or 600×600 pixels. The model trained on images resized to 480×480 achieved a macro precision, recall, and F1-score of 64.3%, 64.3%, and 63%, respectively on the test set. The model trained on images resized to 600×600 pixels achieved a macro precision, recall, and F1-score of 72%, 68.4%, and 66.8%, respectively on the test set. Following these results, 600×600 pixel resolution was selected as the image size for all subsequent trainings. Comprehensive training results can be found in S4 Appendix.

For the best performing model, the validation metrics of the first stage showed an increase in precision from its initial value of 56% to its best value of 90.2% by the end of the first training stage. The validation metrics showed an increase in accuracy from 72% to 87.9% by the end of the second training stage. The macro F1-Score for the best model was 84.2%. After implementing SWA on the top three performing models from the second stage, the model achieved macro precision, recall, and F1-scores of 80.1%, 79.3%, and 79.1%, respectively, on the test set. Compared with the validation metrics, the slight drop-off in F1-score indicates minimal overfitting and the test validation is still within an acceptable range. The model demonstrated classification metrics >= 80% for seven species (Table 2), including four species with more than 100 images and three with fewer than 75 images per class. The model demonstrated classification metrics >= 75% for 10 species.

**Table 2.** Performance metrics of the best performing model on the test subset.

| Species | F1-Score | Precision | Recall |
| --- | --- | --- | --- |
| *Geococcus coffeae* | 97.9 | 100 | 95.8 |
| *Rhizoecus americanus* | 81.5 | 78.6 | 84.6 |
| *Rhizoecus amorphophalli* | 66.7 | 75 | 60 |
| *Rhizoecus cacticans* | 50 | 60 | 42.9 |
| *Rhizoecus coffeae* | 100 | 100 | 100 |
| *Rhizoecus colombiensis* | 57.1 | 57.1 | 57.1 |
| *Rhizoecus dianthi* | 76.2 | 80 | 72.7 |
| *Rhizoecus distinctus* | 88.9 | 100 | 80 |
| *Rhizoecus falcifer* | 96.8 | 100 | 93.8 |
| *Rhizoecus floridanus* | 92.3 | 85.7 | 100 |
| *Rhizoecus leucosomus* | 53.3 | 50 | 57.1 |
| *Ripersiella gracilis* | 66.7 | 60 | 75 |
| *Ripersiella hibisci* | 73.7 | 63.6 | 87.5 |
| *Ripersiella kelloggi* | 97.9 | 100 | 95.8 |
| *Ripersiella kondonis* | 92.3 | 85.7 | 100 |
| *Ripersiella palestineae* | 75 | 85.7 | 66.7 |
| Macro | 79.1 | 80.1 | 79.3 |

*Geococcus coffeae*, *Rhizoecus coffeae*, *R. falcifer*, and *Ripersiella kelloggi* achieved classification metrics >= 90% (Table 2). This followed expectations, as these species contain the most training images in the dataset. These species also exhibit distinctive morphological features (Fig 3), such as the prominent anal lobes and spine-like anal lobe setae of *G. coffeae* or the distinctly rotund body shape of *R. kelloggi*. The model accurately predicted *G. coffeae* with high confidence even when anal lobe setae were absent (Fig 3; CDFA_1316896_2) or were severed from the body and misplaced on the slide (Fig 3; USNMENT02074057_2). The model also correctly identified USNMENT02074057_2 despite it being an immature instar. The model highlighted diffuse areas of *R. kelloggi* for diagnosis but notably did not focus on overlapping folds of the cuticle, which result from flattening rotund specimens, as being important for diagnosis even though this was a common feature of images (Fig 3; USNMENT02075021_2). Instead, it focused on the mouthparts, antennae, medial and marginal regions of the body. *Rhizoecus coffeae* had 100% classification accuracy, which may also be attributed to its rotund body shape in addition to features of the anal ring and mouthparts (Fig 3). For high performing species, Grad-CAM images were relatively consistent, or decisive, regarding the diagnostic features highlighted (Fig 3).

**Fig 3.**
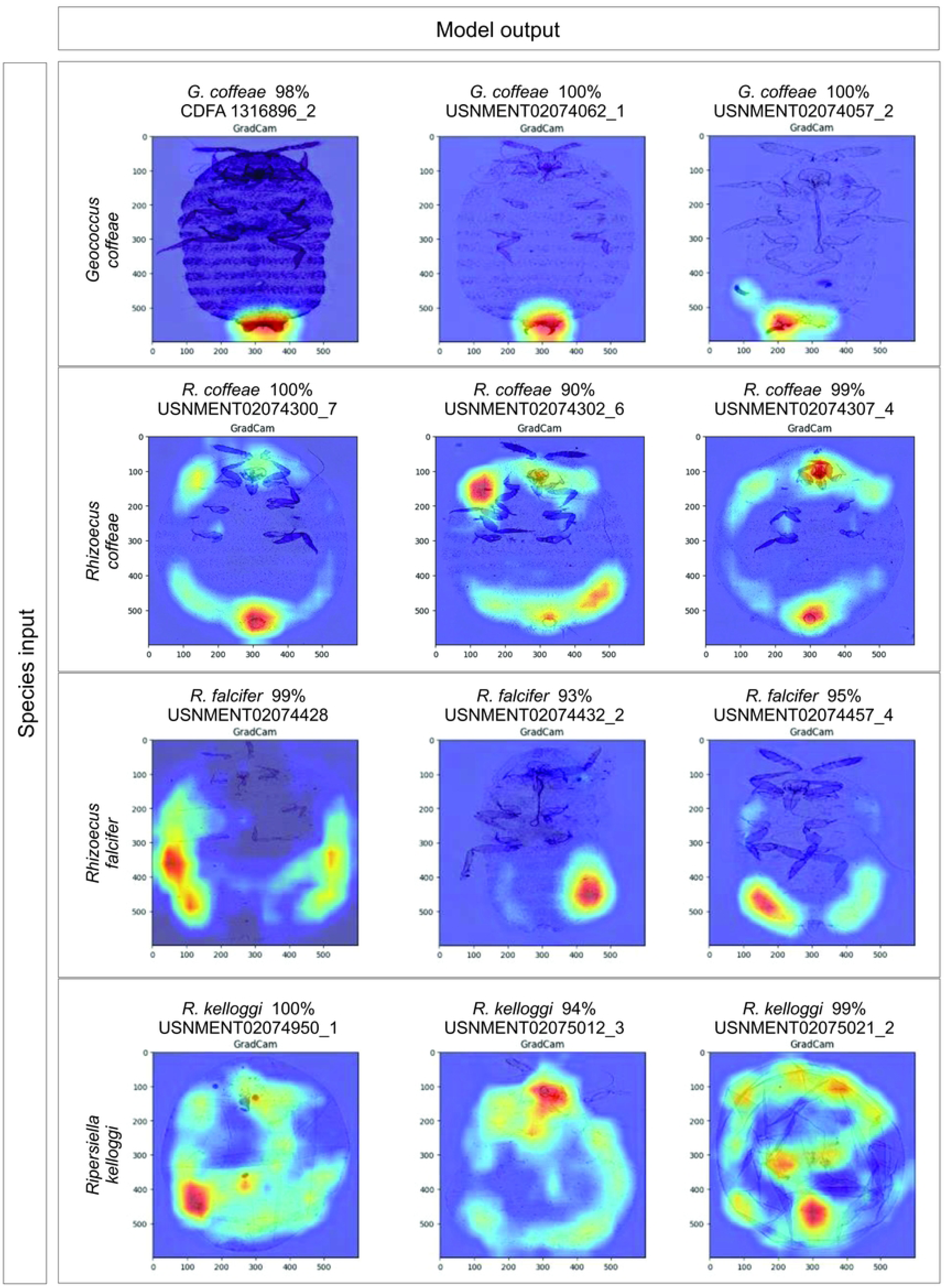
Grad-CAM comparisons for exemplars of the best performing species in the model. Heat maps indicate pixels that most affect final classifications. They point to relevant morphological features and are consistent in identifying traits important for diagnosis.

Certain species performed well even with a limited number of images in the datasets. *Rhizoecus distinctus* contained 47 images, the second-lowest number in the dataset. The species achieved 100% precision and an F1-score and recall at or above 80% (Table 2). Grad-CAM images highlighted the antennae, thoracic margins, anal region, and the distinctly large tubular cerores on abdominal submargins when identifying *R. distinctus* (Fig 4). The model classified one test image from the *R. distinctus* dataset as *Rhizoecus dianthi* (USNMENT02074397_2) (Fig 5). Upon inspection, we discovered that both specimens on the slide are in fact *R. dianthi*. The model correctly identified this specimen, despite having another sample from the slide (USNMENT02074397_1) included in the training set for *R. distinctus*. This shows that the model can be robust to minor human errors, like misidentified specimens or data management errors, when diagnostic features are sufficiently distinctive. *Ripersiella kondonis* also performed well with 57 images, the third-lowest number in the dataset, with a recall of 100% and precision and F1-score at or above 85.7%. Grad-CAM images highlighted the antennae, posterior pair of ostioles, and marginal intersegmental constrictions and/or long marginal setae when identifying *R. kondonis* (Fig 4).

**Fig 4.**
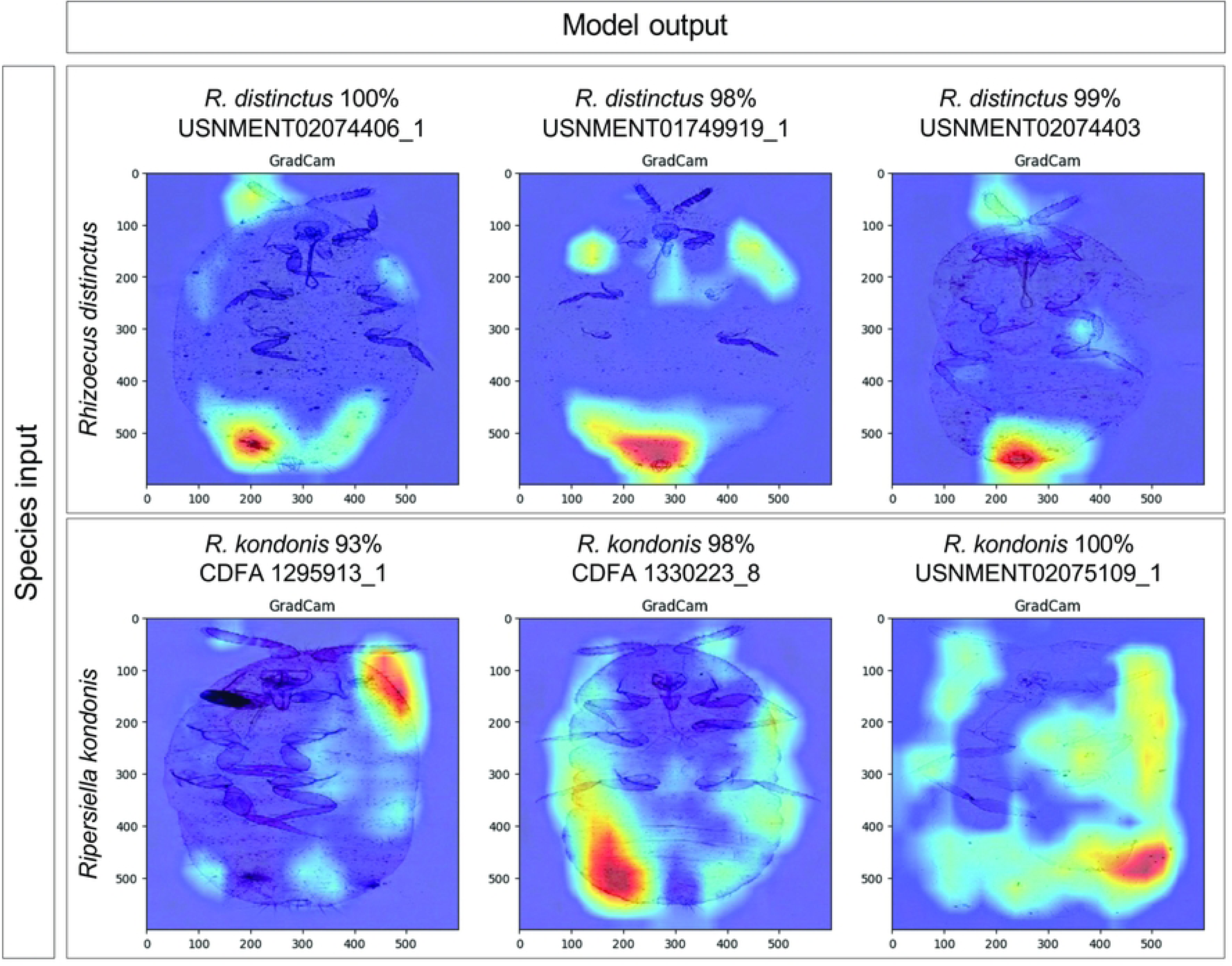
Grad-CAM comparisons for exemplars of high-performing species with distinctive morphological features but relatively few images available for training.

**Fig 5.**
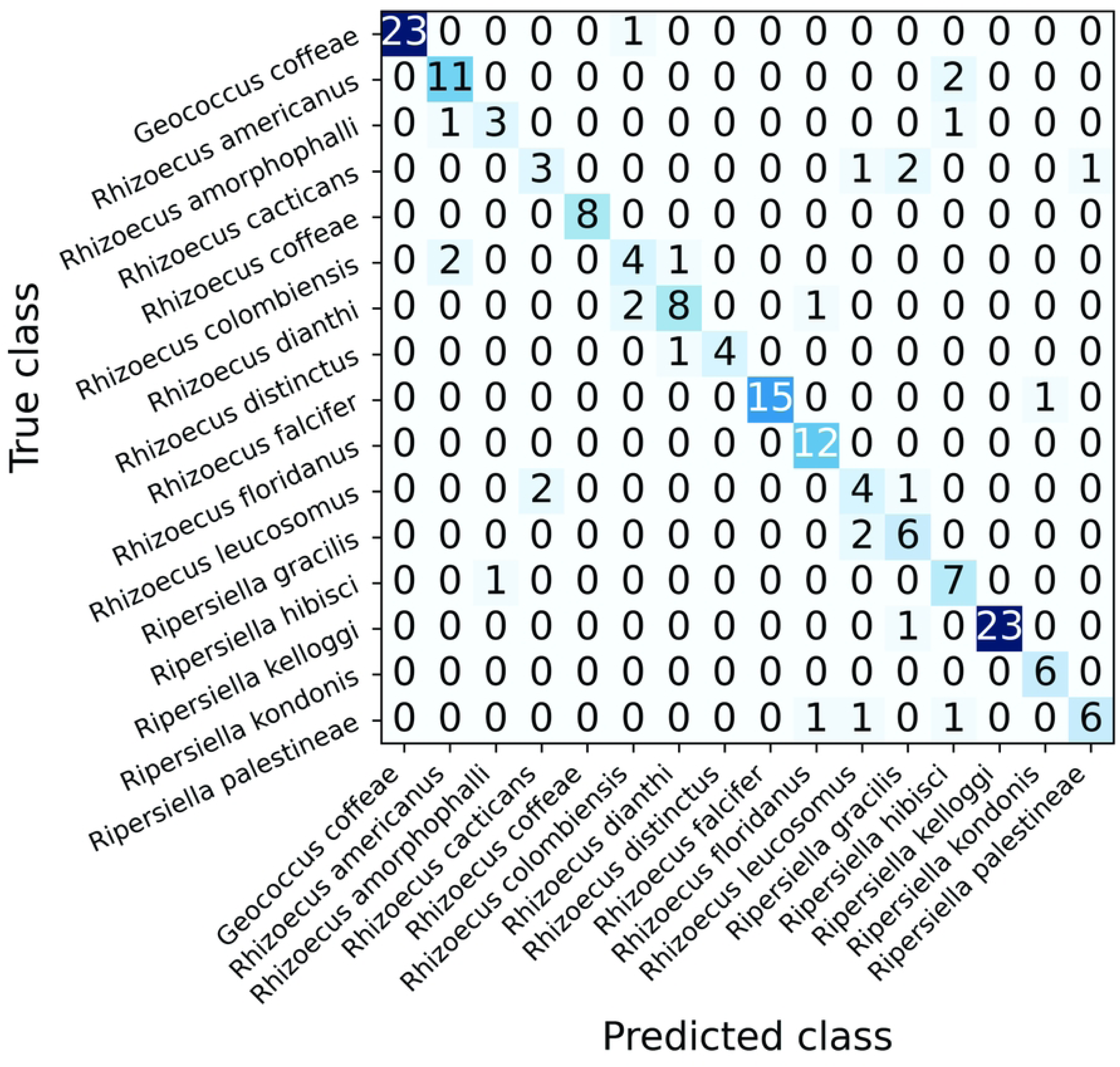
Confusion matrix for the best performing model classifications of the test set. Darker squares represent high correlation between the predicted and true labels.

Underperforming species included *Rhizoecus amorphophalli*, *R. cacticans, R. colombiensis, R. leucosomus*, and *Ripersiella gracilis* (Fig 5), most of which had a limited number of training and test samples, each with fewer than 80 images (Table 1). This behavior was expected, as these species had fewer training images, but may also be attributed to having less distinctive features when compared to species that performed well with few training images, such as *Rhizoecus distinctus* or *Ripersiella kondonis*. For underperforming species, Grad-CAM images appear less decisive in highlighting diagnostic features (Fig 6) compared to high performing species (Fig 3). Focal areas often differed when the model misclassified an image from the test set. Perhaps the model relied more on combinations of traits to correctly predict underperforming species and was more easily misled by variation.

**Fig 6.**
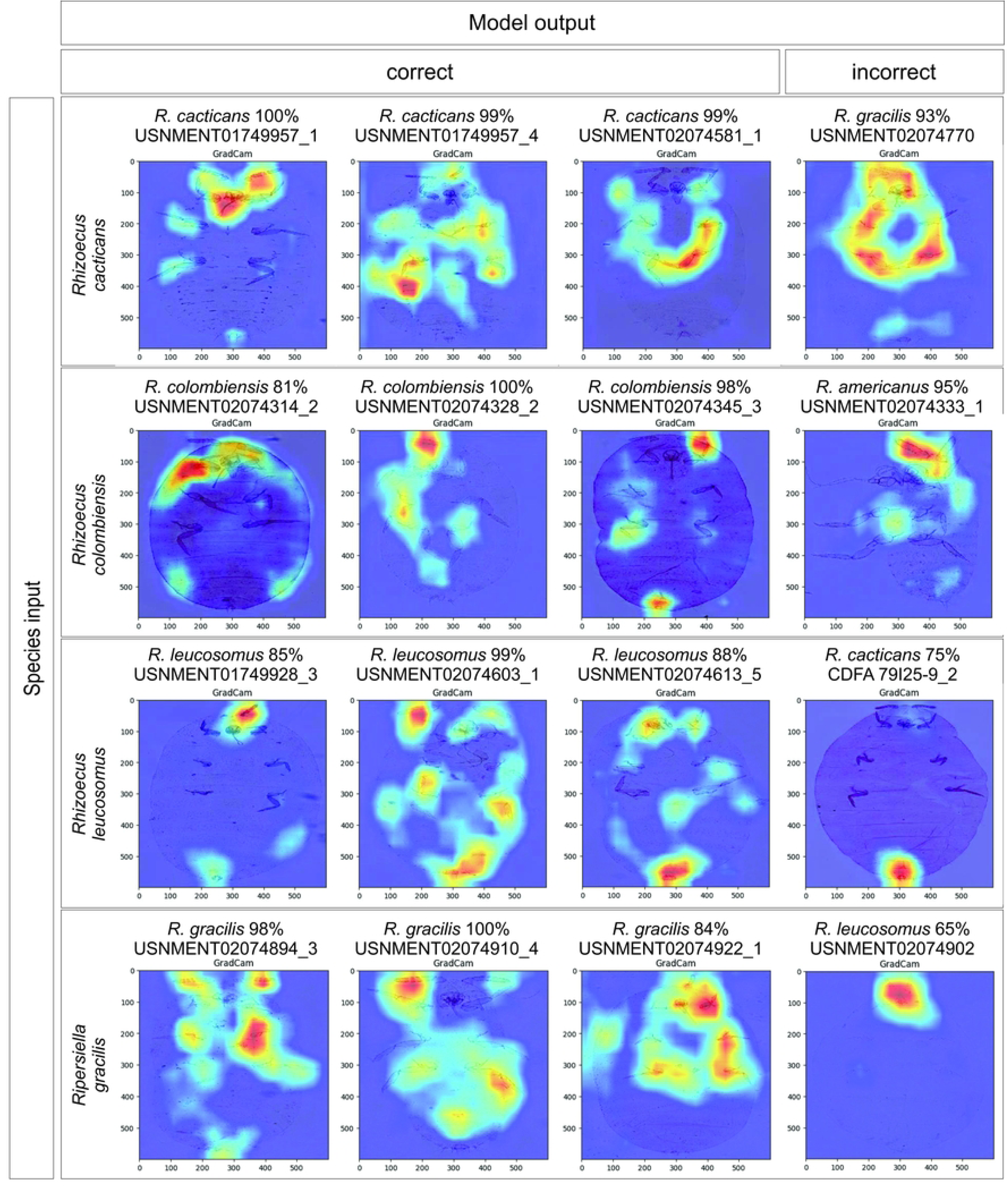
Grad-CAM comparisons for exemplars of underperforming species in the model, including correct and incorrect predictions. Heat maps indicate pixels that most affect final classifications and are variable regarding the features used to diagnose these species.

Misclassification by the model may be attributed, in some cases, to the quality of test images or life stage of specimens but most misidentified specimens were of reasonable quality for identification (S1 Table). Four of 25 misidentified samples (16%) were damaged specimens of poor quality and 3 of 25 (12%) were immature instars; 13 of 25 misidentified samples (52%) were intact, high-quality specimens. All specimens misclassified by the model were re-inspected by the taxonomic expert to confirm their identification. Note that for correct responses from the model, 21 of 144 (15%) were of specimens considered moderately to heavily damaged and 23 of 144 (16%) were immature instars.

### Confidence Scores

Confidence scores differed significantly as a function of prediction outcome (correct vs. incorrect vs. outgroup), Kruskal-Wallis χ^2^ = 64.003, df = 2, p = 1.25e^−14^. Post hoc pairwise Wilcoxon rank-sum tests with Benjamini-Hochberg correction indicated that confidence differed significantly between correct and incorrect responses (p = 1.8e^−11^) and between correct and outgroup responses (p = 3.2e^−7^). But there was no significant difference in confidence between incorrect and outgroup responses (p = 0.11). The correct classifications had significantly higher confidence with a median score of 99% (Table 3), and few responses fell below 80% (Fig 7). Model confidence for incorrect and outgroup classifications were similar, with median scores of 63% and 74%, respectively (Table 3). This suggests that if implemented in a public-facing forum, a classification confidence threshold above 80% would offer a reasonable trade-off, minimizing misclassifications without sacrificing a large proportion of correct predictions that have lower confidence (Fig 7).

**Fig 7.**
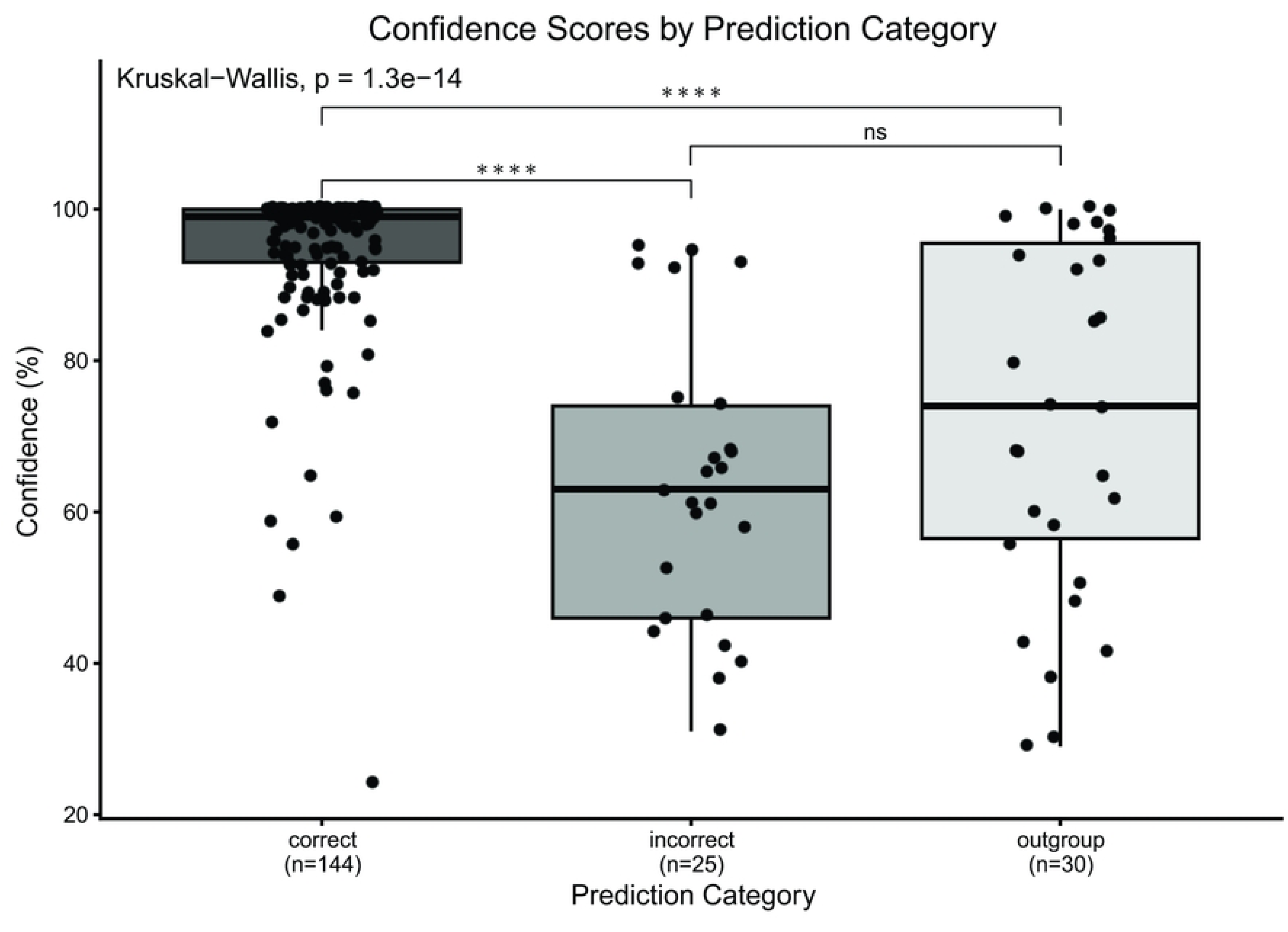
Model confidence by prediction category displayed in a boxplot, significant differences indicated by asterisks. Confidence differs significantly as a function of prediction outcome, Kruskal-Wallis χ^2^ = 64.003, df = 2, p = 1.25e^−14^. Post hoc tests show significant differences between correct and incorrect (p = 1.8e^−11^) and correct and outgroup responses (p = 3.2e^−7^). Incorrect and outgroup responses do not differ significantly (p = 0.11).

**Table 3.**
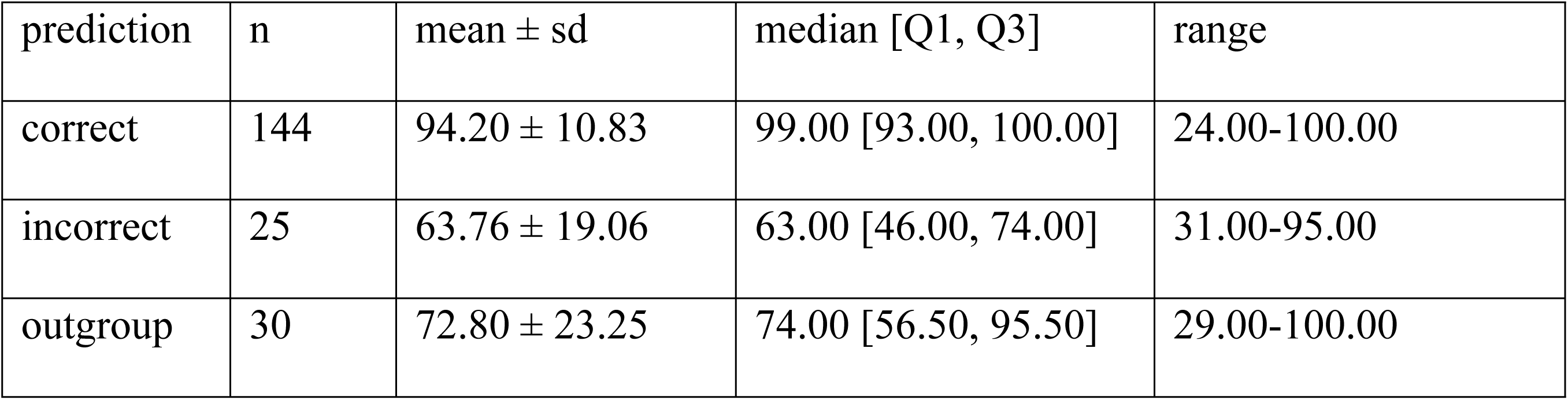
Results of post hoc pairwise Wilcoxon rank-sum tests on confidence scores returned by the model based on correct, incorrect, and outgroup predictions.

## Discussion

This study explored the use of computer vision to identify slide-mounted root mealybugs for rapid identification, with potential downstream applications for regulatory agencies and research. The aim of this study was to lay the foundation for classifying root mealybugs, as an exemplar of slide-mounted insect taxa, using a CNN. The pilot model performed relatively well considering the limited number of images available for some species and an unbalanced dataset typical of natural history collections [37]. The model demonstrated macro-average precision, recall, and F1 scores of 80.1%, 79.3%, and 79.1%, respectively (Table 2). The model demonstrated classification metrics >= 80% for seven species, including some species with fewer than 100 images. The best performing species typically had the largest datasets (Fig 5) and achieved classification metrics >= 90% (Table 2). Yet, some species performed well despite having relatively few images for training, such as *Rhizoecus distinctus* with only 47 images total, indicating that both training set size and distinctiveness of diagnostic features factored into model performance. The worst performing species had relatively fewer images (< 80) and achieved classification metrics between 42–75%.

Grad-CAM images indicate the model used relevant features for species identification, often highlighting traits that taxonomists use in diagnosis, including the antennae, legs, ostioles, anal apparatus, and body shape (Figs 3 and 5). The model recognized relevant diagnostic features of high-performing taxa even on damaged specimens where features were missing or displaced in the image; for example, the anal lobe setae of *Geococcus coffeae* (Fig 3). Additionally, confidence scores assigned to model predictions were significantly higher for correct responses (median = 99%) compared to incorrect responses (median = 63%) (Fig 7; Table 3). When tested against ‘outgroup’ species, root mealybugs that were not included in model training, confidence scores were again significantly lower (median = 74%) in comparison to correct responses (Table 3) and statistically indistinguishable from incorrect responses. This would be a desirable outcome in future real-world applications of a CNN diagnostic tool (Fig 8), where confidence serves as an honest signal and offers useful information about model predictions that require greater scrutiny from a taxonomic expert. Using the current model, a confidence threshold > 80% offers a balanced trade-off that minimizes, but does not entirely preclude, confident misclassifications (Fig 7).

**Fig 8.**
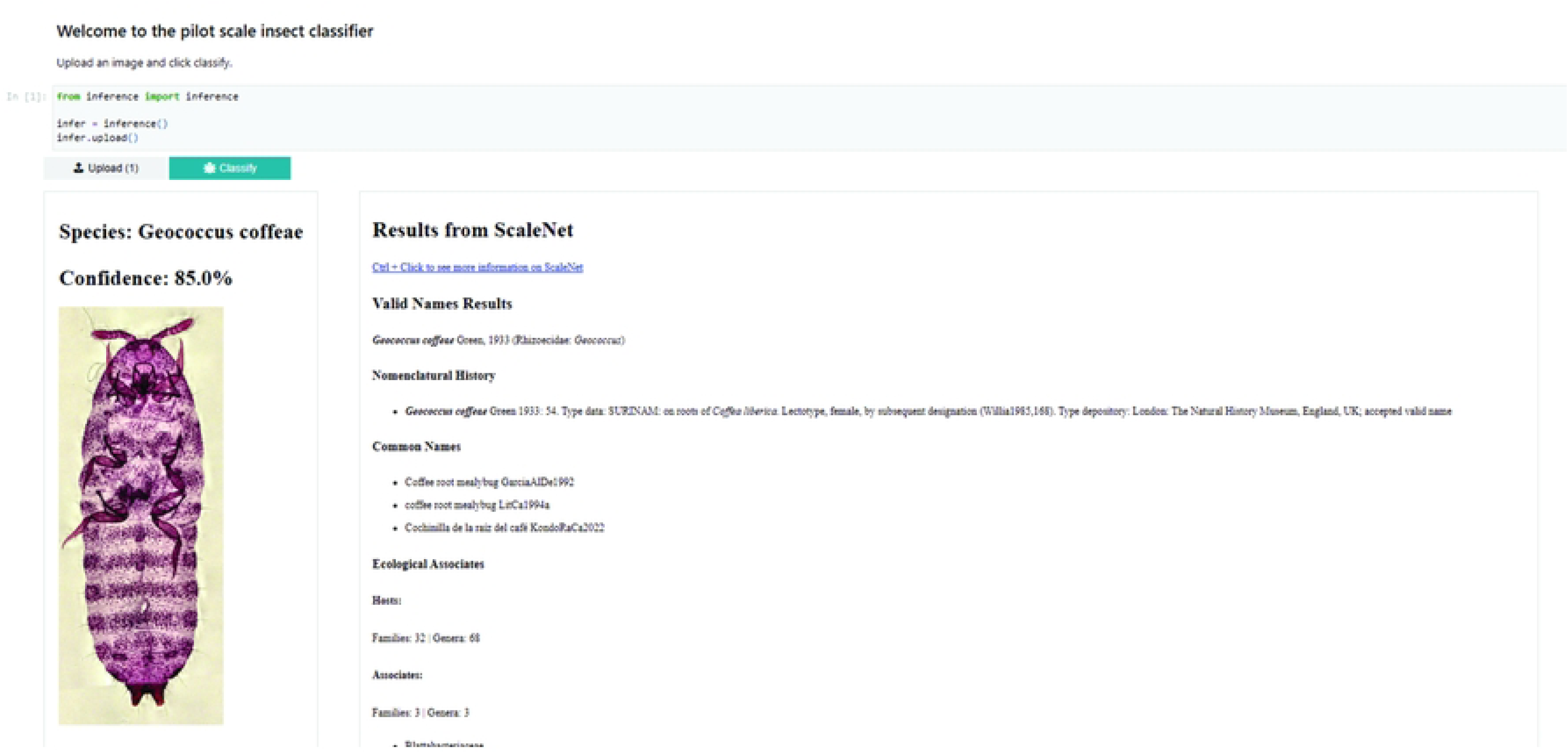
Dashboard of a mock diagnostic CNN tool for the identification of common pests in the family Rhizoecidae. In this example, model output includes the input image, model prediction, confidence score, and species information linked from ScaleNet to support the user in further verification. A pilot version of the dashboard was developed for this study and used in outgroup testing.

Compared to existing studies, our model performance was modest. However, this can be attributed to the lower number of images contained in our study, with the model still showing promise despite this limitation. Many existing species classification applications achieved balanced accuracies greater than 94% [39, 41, 43, 51]. Each of those studies included many training images, typically on the order of 14,000 to 150,000 images; most aimed at a minimum of 200 images per taxa. In the case of Nolte et al. they classified taxa with less than 80 images as underrepresented and combined them into a general ‘other’ class [39]. Simović et al. achieved an overall accuracy of 98.7% for 90 taxa. But in species with fewer images performance was poor, achieving only a 50–75% accuracy, similar to our results for low performers [41]. This indicates that high overall accuracy may be influenced by the number of taxa with more images that achieved a perfect classification rate and is a factor that should be considered in future model development. This is not a limiting factor for future studies, as Motta et al. improved their existing models by increasing the number of training images from 4,056 to 7,561 [38]. They achieved a balanced accuracy of 93.5% for classifying mosquitoes and other insects, an increase from the prior study’s 76.2% [52]. They also employed grid search to optimize the model’s performance. This shows there is potential to improve upon our model by increasing dataset size and further optimizing the model through methods not included in BioEncoder, such as grid or random search, to optimize hyperparameters, which have been shown to improve model performance [52]. Additional considerations include the methodology used for image collection. Recent efforts to digitize natural history collections [50, 53] have expanded the capabilities for automatic taxonomic classification. Yet care must be taken, as whole-drawer imaging can facilitate dataset building, but material may also be misidentified due to human error, unknown cryptic diversity, synonymies, or other taxonomic revisions that are not reflected in a collection. In our small study, with fewer than 2,000 images, there were misidentifications that, if left uncorrected, could have led to poor model performance. For example, we found that some specimens of *R. cacticans* were confused for *R. leucosomus* in the USNM collection, and vice versa. In confirming from these two species were difficult to parse, resulting in the removal of those samples from training out of caution. Based on review of both type series, and in disagreement with some published diagnoses [35], the clearest trait found to distinguish between *R. cacticans* and *R. leucosomus* is the shape of the internal genital organ. The model focused on the genital and anal region to identify *R. leucosomus* to some extent but also emphasized this region when misclassifying a sample as *R. cacticans* (Fig 6; CDFA 79I25-9_2) despite the clear indication of a *leucosomus*-type internal genital organ. Failure of the model to pick up on this distinction may be due to the image resolution used for training, an insufficient number of training images, or a combination of both. This helps identify areas for improvement in future iterations. Note, however, that the model was robust to a real-world example of misidentification for specimens of *Rhizoecus dianthi* that were mislabeled as *R. distinctus* and included in the wrong training and testing datasets. The model highlighted this human error when it correctly identified the test sample (USNMENT02074397_2) as *R. dianthi* with high confidence (99%).

It has been argued for some time that demand for routine identification exceeds what the taxonomic community can accommodate [54]. Demand was expected to rise as early as 2004, and that demand has not let up as local, national, and regional requirements and surveys have grown [54, 55]. The quality of samples needed has always been identified as an issue in terms of quantity, correct identification, standardization (pose, background, and lighting) and rarity of species which continues to be a problem for morphologically similar species [39, 41, 54, 55]. As indicated by the results in this paper, a lack of data needs to be addressed for automatic classifications to perform well. However, the advent of these technologies would support taxonomists in the rapid identification of species encountered in plant quarantine, for example, allowing the community to better accommodate demand for routine identifications and allocate more time to the study of new, cryptic, or otherwise understudied taxa and prioritize systematic research.

To improve performance for use in working environments, we recommend that future research should attempt to increase the dataset, aiming for 200+ images per species ideally. Future studies should incorporate a systematic hyperparameter optimization method and use weighted training to address dataset imbalance for rare species. Additionally, future efforts should address misidentifications that may arise from human error when using a whole-drawer imaging approach. Revisionary systematics research is complementary in this respect; time invested in research improves our understanding of species and their relationships, expands and improves upon natural history collections, and in turn offers more data for model training.

## Acknowledgments

We thank Dr. Erin C. Powell (Florida Department of Agriculture and Consumer Services) and Dr. Natalia von Ellenrieder (California Department of Food and Agriculture) for specimen loans that contributed to this study. This research used resources provided by the SCINet project and/or the AI Center of Excellence of the USDA Agricultural Research Service, ARS project numbers 0201-88888-003-000D and 0201-88888-002-000D. AC was supported by an AI-COE/SCINet Summer Graduate Internship. BR is a participant of the Oak Ridge Institute for Science and Education (ORISE) ARS Research Participation Program, supported by the USDA ARS Systematic Entomology Laboratory in Washington D.C., USA. This research was supported by the U.S. Department of Agriculture, Agricultural Research Service. Mention of trade names or commercial products in this publication is solely for the purpose of providing specific information and does not imply recommendation or endorsement by the U.S. Department of Agriculture; USDA is an equal opportunity provider and employer.

## Supplementary Information captions

S1 Table. Summary of images used in CNN model testing. Includes the species test folder, unique identifiers for images, model prediction and confidence scores, prediction accuracy, group, and additional notes on specimen condition.

S2 Table. Total number of Rhizoecidae images from the USNM, CDFA, and FSCA collections captured with the Glissando slide scanner.

S1 Appendix. Python scripts and a list of all used libraries and their versions. This directory holds all python code and Jupyter Notebooks needed to train, validate, and perform inferencing.

S2 Appendix. Details on training parameters and augmentations. This directory contains the directories generated by BioEncoder natively or that were added during the development process.

S3 Appendix. R code for analysis of model confidence by prediction category. Uses S1 Table as the data matrix.

S4 Appendix. CNN model training results. This directory includes training data, inference, logs, metrics, plots, runs, and weights.

Supplementary information repository: https://doi.org/10.15482/USDA.ADC/33289044

